# Native perennial and non-native annual grasses shape pathogen community composition and disease severity in a California grassland

**DOI:** 10.1101/2020.05.19.104950

**Authors:** Amy E. Kendig, Erin R. Spear, S. Caroline Daws, S. Luke Flory, Erin A. Mordecai

**Affiliations:** Agronomy Department, University of Florida, Gainesville, FL; Smithsonian Tropical Research Institute, Panama City, Panama; Department of Biology, Stanford University, Stanford, CA

**Keywords:** disease severity, pathogen community, host competence, life history, non-native species, grassland, fungi

## Abstract

1. The densities of highly competent plant hosts may shape pathogen community composition and disease severity, altering disease risk and impacts. Life history and evolutionary history influence host competence: longer-lived species tend to be better defended than shorter-lived species and pathogens adapt to infect species with which they have longer evolutionary histories. It is unclear, however, how the densities of species that differ in competence due to life and evolutionary histories affect plant pathogen community composition and disease severity.
2. We examined foliar fungal pathogens on two host groups in a California grassland: native perennial and non-native annual grasses. We first characterized pathogen community composition and disease severity on the two host groups to approximate differences in competence. We then used observational and manipulated gradients of native perennial and non-native annual grass densities to assess the effects of each host group on pathogen community composition and disease severity in 1-m^2^ plots.
3. Native perennial and non-native annual grasses hosted distinct pathogen communities but shared generalist pathogens. Native perennial grasses hosted pathogens with larger host ranges and experienced greater disease severity (75% higher proportions of leaves with lesions) than non-native annuals. While the relative abundances of three common pathogens tended to shift, and disease severity of native perennial grasses tended to increase with increasing densities of both host groups, these changes were not statistically significant.
4. *Synthesis.* The life and evolutionary histories of grasses likely influence their competence for different pathogen species, leading to distinct pathogen communities and differences in disease severity. However, there was no evidence that the density of either host group significantly affected pathogen community composition or disease severity. Therefore, variation in competence for different pathogens likely shapes pathogen community composition and disease severity but may not interact with host density to alter disease risk and impacts at small scales.

## Introduction

Community composition can affect infectious disease risk and impacts (Rohr et al., 2020; Zhu et al., 2000). Organisms serve as habitat patches for pathogens and other microbes (Borer, Laine, & Seabloom, 2016; Mihaljevic, 2012) and, synonymous with habitat quality and area, the competence and density of hosts are expected to affect pathogen persistence and prevalence (Burdon & Chilvers, 1982; Fenton, Streicker, Petchey, & Pedersen, 2015). Communities with higher densities of competent hosts (i.e., those with high susceptibility and/or ability to transmit pathogens) should experience greater disease risk and impacts (Joseph, Mihaljevic, Orlofske, & Paull, 2013; Young, Parker, Gilbert, Sofia Guerra, & Nunn, 2017). Because the relationship between community composition and disease can impact biodiversity conservation (Rohr et al., 2020), empirical studies of natural communities tend to focus on species richness more than host density (but see Mitchell, Tilman, & Groth, 2002; Schmidt et al., 2020; Wojdak, Edman, Wyderko, Zemmer, & Belden, 2014). It is therefore unclear how the densities of hosts that differ in competence drive disease risk and impacts.

Infection prevalence (i.e., proportion of hosts infected) and disease severity (e.g., proportion of leaf area with fungal lesions) are indicators of disease risk (Rohr et al., 2020). Typically, multiple pathogens circulate within host communities (Dobson, Lafferty, Kuris, Hechinger, & Jetz, 2008; Griffiths, Pedersen, Fenton, & Petchey, 2011; Malpica, Sacristán, Fraile, & García-Arenal, 2006), driving infection prevalence and disease severity (Alizon, de Roode, & Michalakis, 2013; Mordecai, Hindenlang, & Mitchell, 2015; Vasco, Wearing, & Rohani, 2007). Variation among hosts in competence for different pathogens, which can arise through variation in traits and evolutionary histories with pathogens (Barrett, Kniskern, Bodenhausen, Zhang, & Bergelson, 2009; Joseph et al., 2013; I. M. Parker & Gilbert, 2004), can promote diversity in pathogen communities (Johnson et al., 2016). Therefore, variation in life history and evolutionary history may alter disease risk through pathogen community composition.

Life history and evolutionary history can influence plant–pathogen interactions and may affect pathogen community composition and overall disease severity. Shorter-lived species, such as annual plants, are expected to be less well-defended against pathogens and experience greater disease severity than longer-lived species because they allocate more resources to reproduction than survival (Cronin, Welsh, Dekkers, Abercrombie, & Mitchell, 2010; Joseph et al., 2013; Miller, White, & Boots, 2007). Plant species with longer evolutionary histories with pathogens in a given location may be more susceptible to attack by specialists that have overcome specific plant defenses (I. M. Parker & Gilbert, 2004; Telfer & Bown, 2012). In addition, species introduced to a new geographic area are likely to leave their specialist pathogens behind, as predicted by the enemy release hypothesis (Keane & Crawley, 2002). However, non-native plants tend to be annual species (Razanajatovo et al., 2016; Sutherland, 2004) and can accumulate pathogens over time (Hawkes, 2007), suggesting that long-established non-native plants may have overlapping pathogen communities with native species.

In general, higher densities of plant assemblages result in more contacts between susceptible host tissue and pathogen propagules (Burdon & Chilvers, 1982) and increase the negative effects of infection (Lively, Johnson, Delph, & Clay, 1995). Changes in the density of a single plant genotype or species are more likely to affect specialist pathogens than generalist pathogens (Alexander & Holt, 1998), but specific plant traits may interact with plant density to promote infection by generalist pathogens. For example, non-native annual grasses in California grasslands may increase pathogen transmission by filling in the gaps between native perennial bunchgrasses (S. S. Parker, Seabloom, & Schimel, 2012) and native perennial grasses may grow later into the growing season than non-native annuals (Chiariello, 1989; Seabloom, Harpole, Reichman, & Tilman, 2003), providing additional opportunities for pathogen infection and dispersal (Thrall, Biere, & Antonovics, 1993). Thus, pathogen communities may shift, and disease severity may increase with increasing density of either non-native annuals or native perennials, but to a greater extent with increasing density of the more competent group.

Here we assess how the densities of native perennial and non-native annual grasses affect foliar fungal pathogen community composition and disease severity in a California grassland. California grasslands are dominated by non-native annual grasses, which differ in life history and local residence time from native perennial bunchgrasses (Heady, 1977). Non-native annual grasses have been established in California for more than a century (Heady, 1977) and serve as hosts for a diversity of foliar fungal pathogens that are transmitted through density-dependent mechanisms (McCartney, Fitt, & West, 2006; Spear & Mordecai, 2018). We first approximated differences in competence by answering the question: *How do (a) pathogen community composition and (b) disease severity differ between native perennial and non-native annual grass hosts?* Pathogen community composition and disease severity depend on, among other factors, multiple aspects of host competence: susceptibility, resistance, tolerance, and transmission (Barrett et al., 2009; Borer et al., 2016; Susi, Vale, & Laine, 2015). We hypothesized that native perennials would host more specialized pathogens due to longer evolutionary history with local pathogens, that non-native annuals would experience higher disease severity due to lower allocation to defenses, and that the two groups would host overlapping pathogen communities due to the long residence time of non-native annuals. We then evaluated the effects of host density on disease risk by answering the question: *How do native perennial and non-native annual grass densities affect (a) pathogen community composition and (b) disease severity?* We hypothesized that increasing densities of either non-native annuals or native perennials would shift pathogen communities and increase disease severity, and that the density of the more competent group would have a larger effect.

## Materials and methods

### Study system

We carried out foliar fungal sampling and quantified disease severity from 2015 to 2017 at Stanford University’ s Jasper Ridge Biological Preserve (JRBP) in San Mateo County, California, USA. California grasslands are dominated by non-native Mediterranean annual grasses that rapidly established during European settlement (Bossard & Randall, 2007; Hamilton, 1997). The non-native annual grass species included in this study were: *Avena barbata, Avena fatua, Brachypodium distachyon, Bromus diandrus, Bromus hordeaceus, Bromus sterilis, Festuca myuros*, and *Gastridium phleoides*. All of these species except *B. sterilis* and *G. phleoides* are categorized as invasive in California (Cal-IPC, 2020). *Avena* and *Bromus* species dominate the non-serpentine grasslands at JRBP (McNaughton, 1968). *Stipa pulchra*— hypothesized to be the dominant California grassland species prior to European settlement (Heady, 1977)—and *Elymus glaucus* are the two native perennial grasses included in this study. A study at JRBP in 2015 demonstrated that unique pathogen communities were associated with several grass species, but that generalist pathogens were shared among grass species (Spear & Mordecai, 2018). The data from that study are included here, along with additional data collected in the two following years. Plant growth at JRBP begins with the onset of precipitation in the late fall, progresses through the cool, wet winters into the spring, and ends in the warm, dry summers (Chiariello, 1989). The cumulative precipitation in San Mateo county between September and April was 579 mm (2014-2015), 728 mm (2015-2016), and 1139 mm (2016-2017), ranging on both sides of the 100-year average of 683 mm (NOAA, 2020).

### Density gradients

In the spring of 2015, we established ten observational transects across visually-assessed gradients of perennial grass–dominated to annual grass–dominated areas of JRBP (“observational study”; Spear & Mordecai, 2018; Uricchio, Daws, Spear, & Mordecai, 2019). Transects consisted of four to five 1-m^2^ plots (Fig. S1) and were sampled over two years (2015 and 2016). Following the first sampling, we supplemented transects that lacked common non-native annual or native perennial species by planting seven background plant groups— approximately 20 seeds each of *A. barbata, B. diandrus, B. hordeaceus, E. glaucus*, and *S. pulchra* and adult *E. glaucus* and *S. pulchra* transplanted from elsewhere in JRBP—into the first, middle, and last plots. We counted the number of individuals per grass species within subplots, and scaled the counts up to 1-m^2^, for 47 and 18 plots during April 8–9, 2015 and June 28–July 5, 2016, respectively (Table S1). We assumed that forbs were outside of the primary host range of the focal fungal pathogen species (Gilbert & Webb, 2007; I. M. Parker et al., 2015) and therefore did not include forb density in our analyses even though forbs were present in the plots.

In the fall of 2015, we established 70 1-m^2^ plots in a 35 m × 35 m area of JRBP where weed matting had been placed for several months to suppress background plant recruitment (“manipulated experiment”, Fig. S1; Uricchio et al., 2019). Within the 1-m^2^ plots, we manipulated the densities of the seven background plant groups to 10%, 20%, 40%, 80%, or 100% of the density of each in monoculture by sowing seeds and transplanting adult plants (two replicate plots per treatment). In addition, ten 2 x 2 m plots were cleared and planted with one individual from each background plant group. In January 2016, we added “focal” individuals to the plots by sowing ten seeds or one adult from each background plant group. During June 1–24, 2016, we counted up to 50 individual grasses in each plot, identified them to species, and scaled the densities to 1-m^2^ (Table S1). We weeded species that were not planted throughout the growing season, but some survived. We included the densities of unintentional grasses in our analyses along with intentional plants.

In both the observational study and manipulated experiment, the grass communities included either high non-native annual grass density and low native perennial grass density, high native perennial grass density and low non-native annual grass density, or low densities of both (Fig. S2A–B). The majority of the plots had more non-native annual grasses than native perennial grasses, with native perennial grass relative abundance reaching a maximum of about 0.5 in the observational study and 0.8 in the manipulated experiment (Fig. S2C–D).

### Foliar fungal pathogen communities

To evaluate the differences between foliar fungal pathogen communities associated with native perennial and non-native annual grasses, we collected leaf tissue with lesions from plants within the observational study, the focal plants in the manipulated experiment, and from additional plants located across JRBP (Fig. S1). Between March and June each year, we obtained isolates from the foliar fungal lesions of 91, 261, and 182 native perennial grasses and 75, 242, and 110 non-native annual grasses, in 2015, 2016, and 2017, respectively. One piece of leaf tissue was collected per plant. Native perennials included *E. glaucus* and *S. pulchra* and non-native annuals included *A. barbata, A. fatua, B. diandrus*, and *B. hordeaceus*.

As described by Spear and Mordecai (2018), we isolated fungi associated with lesions by excising symptomatic tissue from the edge of foliar lesions and surface sterilizing (sequential immersion in 70% ethanol and 10% household bleach, 60 s each) and plating each tissue piece on 2% malt extract agar (MEA) with chloramphenicol. For each tissue piece, morphologically distinct hyphae (i.e., morphotypes) were isolated into pure culture on 2% MEA plates. The Mordecai lab maintains reference strains (California Department of Food and Agriculture permit 3160). For each fungal isolate, we extracted genomic DNA and amplified the internal transcribed spacer (ITS) regions 1 and 2, the 5.8S rRNA gene, and part of the rRNA LSU as detailed in Spear and Mordecai (2018). However, in 2017, we modified our protocol to produce longer consensus reads. Specifically, we paired the forward primer ITS1-F (Gardes & Bruns, 1993) with the reverse primer LR3 (Vilgalys & Hester, 1990), rather than ITS4-B (Gardes & Bruns, 1993).

We processed the Sanger sequencing reads from MCLAB (San Francisco, California, USA) with Geneious 7.1.9 (Kearse et al., 2012). We trimmed and automatically assembled reads when possible; when not possible, we manually assembled reads or selected the longest trimmed individual read over 100 bp. We clustered all consensus sequences into operational taxonomic units (OTUs) based on 97% sequence similarity using USEARCH 10.0.240 (Edgar, 2010). If different morphotypes from the same tissue piece were clustered into the same OTU, we assumed they represented the same isolate. We estimated the taxonomic placement of the ITS OTUs with the UNITE database 01.12.2017 (Nilsson et al., 2019) and assigned taxonomy in mothur 1.40.5 (Schloss et al., 2009) using the naïve Bayesian classifier (Wang, Garrity, Tiedje, & Cole, 2007) with a bootstrapping confidence score of at least 80% for species name and at least 60% for any other taxonomic rank.

### Disease severity

We assessed disease severity caused by foliar fungi by haphazardly selecting one to six leaves, based on availability, per plant and visually approximating the proportion of leaf area covered by lesions. We assessed leaves from native perennial and non-native annual grasses with the following sample sizes (in this order): 101 native perennial and 292 non-native annual grasses in the observational study during March 11–16, 2015; 71 and 129 grasses in the observational study and along four additional transects at JRBP (T11-T14, Fig. S1) during April 17–20, 2015; 69 and 11 grasses in the observational study during May 25–27, 2016; and 269 and 161 grasses in the manipulated experiment during May 5–24, 2016. The sampling in March and April of 2015 used the same marked plants, so we analyzed these data separately. We present results with March data in the main text and provide results with April data in the supplementary materials for comparison. Native perennials included *E. glaucus* and *S. pulchra* and non-native annuals included *A. barbata, B. diandrus*, and *B. hordeaceus*.

### How does pathogen community composition differ between native perennial and non-native annual grass hosts?

To evaluate differences between the foliar fungal communities associated with each host group, we performed a permutational multivariate analysis of variance (PERMANOVA) using the Chao method to calculate community dissimilarity. The Chao method accounts for unobserved species and is robust to differences in sample sizes (Chao, Chazdon, & Shen, 2005). We defined a community as the foliar fungal isolates associated with a grass species in a particular year (Table S2) and created a community matrix of isolate abundance with each community as a row and each OTU as a column. We omitted communities with fewer than four isolates, leading to six communities associated with native perennials and ten communities associated with non-native annuals. We used the community matrix as the response variable in the PERMANOVA and the grass species, year, and host group as the predictor variables. We used non-metric multidimensional scaling (NMDS) with the Chao method to visualize differences among foliar fungal communities.

We also estimated the known host ranges of pathogens associated with each host group to evaluate whether differences in pathogen community composition can be at least partially explained by escape of non-native annual grasses from specialist pathogens (Keane & Crawley, 2002). We searched the U.S. National Fungus Collections Database (Farr & Rossman, 2019) for the number of host species associated with the OTUs for which we estimated a species name (Schmidt et al., 2020). To test the hypothesis that the pathogens associated with the two host groups differ in their host ranges, we performed a Welch two sample t-test, comparing the host ranges of fungal isolates associated with native perennial grasses to those of fungal isolates associated with non-native annual grasses. By using each fungal isolate as a replicate, estimated fungal species that were isolated more frequently contributed more to the average host range. Note that the database may provide more information, and potentially larger host range estimates, for fungi of crops and economically important plants, fungi intercepted at ports of entry, common fungi, and invasive or emerging fungal pathogens (Farr & Rossman, 2019).

### How does disease severity differ between native perennial and non-native annual grass hosts?

To evaluate the differences in disease severity between native perennial and non-native annual grasses, we used two metrics for each plant: the proportion of leaves with lesions (“leaves”) and, for the leaves with lesions, the proportion of leaf surface area covered (“surface”). The “leaves” metric approximates the exposure of each leaf to fungal spores and susceptibility to tissue colonization by pathogens. The “surface” metric approximates the plant’ s ability to suppress fungal growth in the tissue (i.e., resistance). We fit a generalized linear mixed effect model with a logit-link binomial error distribution to the presence/absence of lesions for each leaf for the “leaves” metric and a generalized linear mixed effect model with a logit-link beta error distribution to proportion of leaf surface area with lesions for each infected leaf for the “surface” metric. The predictor variable was whether or not the plant was a non-native annual and the random effect intercepts were the plant ID nested within plot nested within experiment and crossed with year. We removed experiment from the random effects for the “surface” model (variance < 8×10^−18^) to help with model convergence; spatial heterogeneity was still accounted for with the random effect “plot”.

### How do native perennial and non-native annual grass densities affect pathogen community composition?

To assess the effects of native perennial and non-native annual grass densities on the composition of foliar fungal pathogen communities associated with each of these groups, we evaluated the change in isolation frequency of the most common OTUs over the density gradients. We used a subset of the full fungal isolate dataset, selecting only those isolates collected from the density gradients. Sample sizes in the order of native perennial and non-native annual grasses were: 69 and 55 isolates from 31 plots of the observational study in 2015, 22 and 17 isolates from six plots of the observational study in 2016, and 135 and 76 isolates from the 28 plots of the manipulated experiment planted at 80% and 100% density. Native perennials included *E. glaucus* and *S. pulchra*; non-native annuals included *A. fatua* (observational study only), *A. barbata, B. diandrus*, and *B. hordeaceus*.

To select the most common OTUs, we ranked all of the OTUs from the full fungal isolate dataset by the number of isolates obtained in each year (i.e., abundance). We evaluated the differences in abundance between consecutive ranks and found relatively large differences between the fifth and sixth most common OTUs in 2015 and 2016 and between the fourth and fifth most common OTUs in 2017 (Fig. S3). Therefore, we selected the top five, five, and four most common OTUs in 2015, 2016, and 2017, respectively, which resulted in seven focal OTUs. The fungal species associated with these OTUs were *Alternaria infectoria, Parastagonospora avenae, Pyrenophora chaetomioides, Pyrenophora lolii, Pyrenophora tritici-repentis*, an unidentified *Pyrenophora* species, and *Ramularia proteae*. Note that we refer to the OTUs by their estimated species names in the results, but that these same species names may be associated with less common OTUs as well.

We fit generalized linear mixed effect models with logit-link binomial error distributions to presence/absence of each focal OTU, analyzing the observational study and manipulated experiment separately. The presence/absence data indicate whether a foliar fungal isolate was identified as the OTU of interest. The fixed effect predictor variables were the host group from which the fungal isolate was collected, native perennial grass density and non-native annual grass density in the plot from which the fungal isolate was collected, and, when present, the density of grasses that were either unidentified or not included in either of the groups (“other grasses”, Table S1). The fixed effects also included interactions between the host group and each of the grass density measurements. Random effect intercepts included the plot crossed with year for the observational study and the plot for the manipulated experiment. We performed model selection by fitting all fixed effect subset models for each global model. The Akaike information criterion with a correction for small sample sizes (AICc) was calculated for each model and we extracted the subset of the models for which the cumulative sum of the normalized model likelihoods was greater than or equal to 0.95 (i.e., the 95% confidence set of models). We report coefficient estimates from the average of the 95% confidence set.

Among the set of the most common fungal species, models were not fit for *P. chaetomioides* on native perennial grasses, *P. tritici-repentis* on non-native annual grasses, and *R. proteae* in the manipulated experiment because of low sample sizes (in all cases, *n* < 2 isolates). To allow model convergence, we removed the interaction between host group and “other grass” density for *P. lolii* and *R. proteae* in the observational study, between host group and all types of grass density for the unidentified *Pyrenophora* species and *P. avenae* in the observational study, and between host group and both types of grass density for *P. lolii* in the manipulated experiment.

### How do native perennial and non-native annual grass densities affect disease severity?

To evaluate the effects of native perennial and non-native annual grass densities on the disease severity of each host groups, we used the same metrics already described: “leaves” and “surface”. We used a subset of all of the disease severity assessments, selecting only those collected from the density gradients. Sample sizes in the order of native perennial and non-native annual grasses were: 101 and 292 grasses from 46 plots of the observational study in March 2015, 56 and 79 grasses from 25 plots of the observational study in April 2015, 69 and 11 grasses from 18 plots of the observational study in 2016, and 269 and 161 grasses from all plots of the manipulated experiment. Native perennials included *E. glaucus* and *S. pulchra*; non-native annuals included *A. barbata, B. diandrus*, and *B. hordeaceus*.

Again, we fit generalized linear mixed effect models with logit-link binomial (for “leaves”) and beta (for “surface”) error distributions to presence/absence of lesions for each leaf (for “leaves”) and proportion of leaf surface area with lesions for each infected leaf (for “surface”). The fixed effects were consistent with those described for the pathogen isolate models in the preceding section. The random effect intercepts were the plant ID nested within plot and crossed with year for the observational study and plant ID nested within plot for the manipulated experiment. We also performed model selection as described in the preceding section, except for the observational study “surface” model—sub-models could not converge during model averaging. Statistical analyses were completed in R version 3.5.2 (R Core Team, 2018) using vegan (Oksanen et al., 2019), rusda (Krah et al., 2018), glmmTMB (Brooks et al., 2017), MuMIN (Bartoń, 2019), DHARMa (Hartig, 2019), and tidyverse (Wickham, 2017).

## Results

### How does pathogen community composition differ between native perennial and non-native annual grass hosts?

We identified 83 unique OTUs from the 961 foliar fungal isolates collected from six grass species at JRBP (Fig. 1A). Forty-one OTUs were isolated from only native perennial grasses, 18 were isolated from only non-native annual grasses, and 24 were isolated from both host groups. The host groups explained 25% of the variance in pathogen community composition (Table 1) and the pathogen communities associated with the two groups were distinct (Fig. 1B). However, the 24 OTUs isolated from both host groups made up 78% and 96% of the isolates from native perennial grasses and non-native annual grasses, respectively, leading to overlap between the pathogen communities associated with the two host groups (Fig. 1A). Fungal species names (29 total) were estimated for 282 native perennial grass isolates (53%) and 266 non-native annual grass isolates (62%). The estimated fungal species isolated from non-native annual grasses had, on average, smaller host ranges than those isolated from native perennial grasses (Fig. 1C, Welch two sample t-test: t = 4.53, df = 480, *P* < 0.001).

**Table 1.** PERMANOVA describing the effects of host group, grass species, and sampling year on pathogen community composition.

|  | df | Sums of sqs. | Mean sqs. | F | R<sup>2</sup> | *P* |
| --- | --- | --- | --- | --- | --- | --- |
| Host group | 1 | 0.516 | 0.516 | 23.506 | 0.250 | **0.001** |
| Grass species | 4 | 0.510 | 0.127 | 5.801 | 0.247 | **0.004** |
| Year | 2 | 0.862 | 0.431 | 19.606 | 0.417 | **0.001** |
| Residuals | 8 | 0.176 | 0.022 | NA | 0.085 | NA |
| Total | 15 | 2.064 | NA | NA | 1 | NA |

df = degrees of freedom, sqs. = squares, NA = not applicable; *P*-values indicating statistical significance ( $P < 0.05$ ) are in bold.

**Figure 1.**
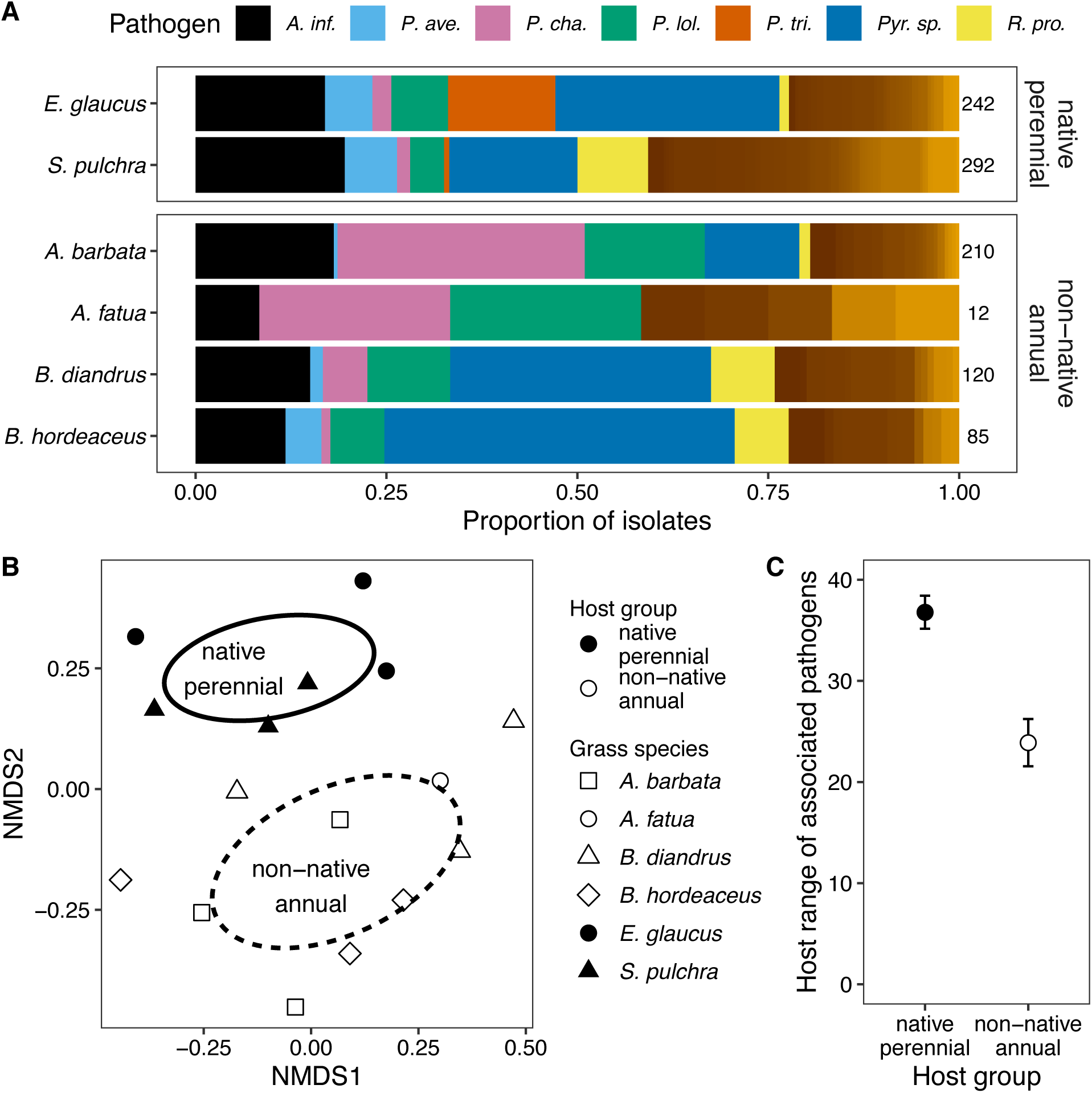
Pathogen communities associated with native perennial and non-native annual grasses. (A) Composition of fungal pathogen isolates for each grass species. Each OTU is represented by a different color and the legend is provided for the seven focal OTUs: *Alternaria infectoria* (*A. inf.*), *Parastagonospora avenae* (*P. ave.*), *Pyrenophora chaetomioides* (*P. cha.*), *Pyrenophora lolii* (*P. lol.*), *Pyrenophora tritici-repentis* (*P. tri.*), *Pyrenophora sp.* (*Pyr. sp.*), and *Ramularia proteae* (*R. pro.*). The total number of isolates per grass species is to the right of the bars. (B) NMDS plot of pathogen communities associated with the two host groups. A “community” is all of the foliar fungal isolates from one grass species in a year. (C) The average number of host species in the USDA Fungal Database for fungal pathogens isolated from each host group (mean ± 1SE). Averages are composed of all fungal isolates with estimated species names and available data; the host ranges of pathogens more frequently isolated contributed proportionally more to the summary statistics.

### How does disease severity differ between native perennial and non-native annual grass hosts?

Native perennial grasses had 75% higher proportion of leaves with lesions than non-native annual grasses (Fig. 2A, *P <* 0.01, Table S3). The proportion of leaf area with lesions was generally low and did not differ between native perennial and non-native annual grasses (Fig. 2B, *P* = 0.93, Table S3). These patterns were maintained when data collected in April 2015 from the observational study were substituted for data collected in March 2015 (Fig. S4).

**Figure 2.**
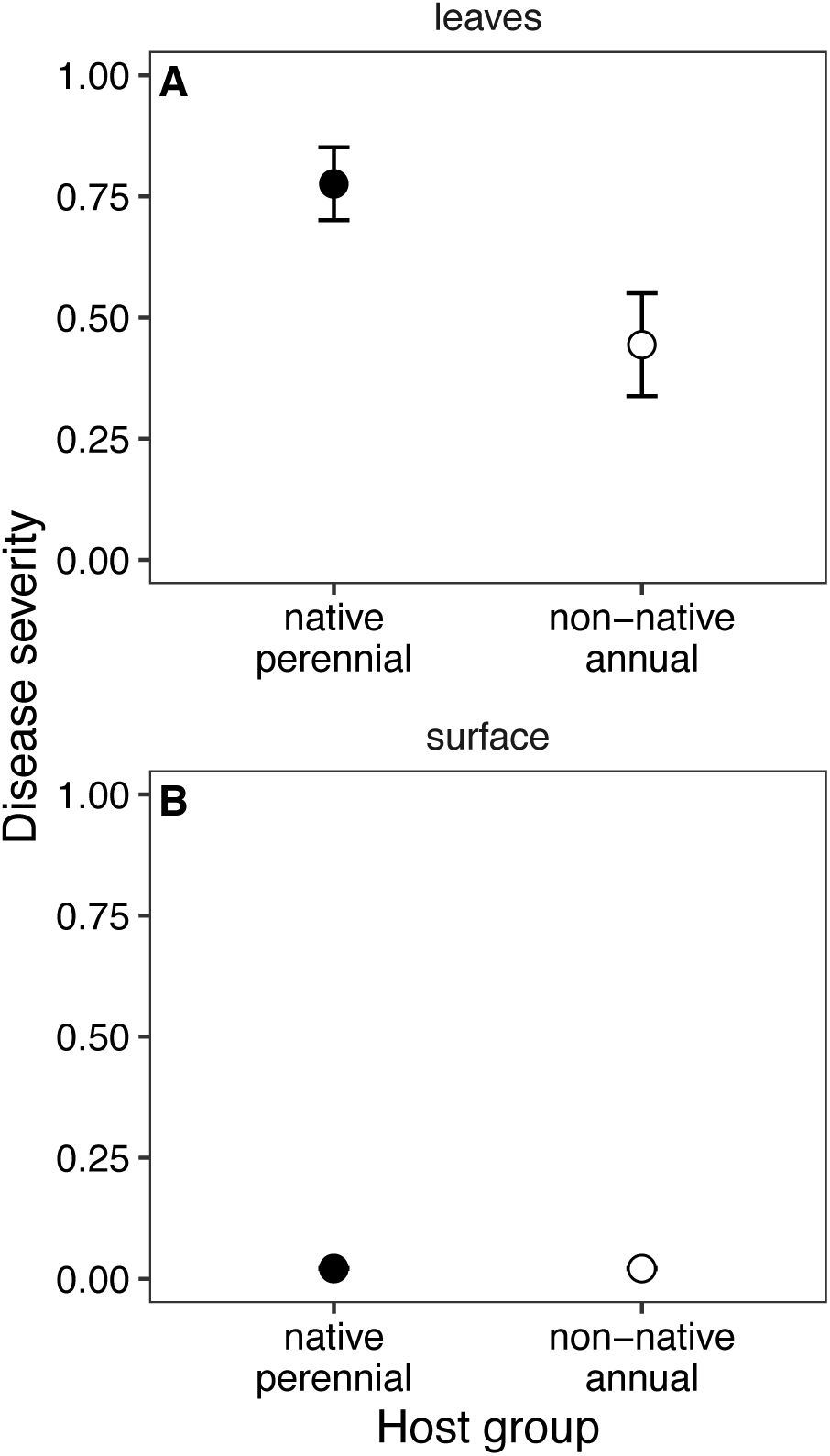
The average disease severity for the native perennial and non-native annual host groups (model-estimated mean ± 1SE), quantified as (A) the proportion of leaves with lesions per plant (“leaves”) and (B) the average proportion of infected leaf surface area with lesions per plant (“surface”).

### How do native perennial and non-native annual grass densities affect pathogen community composition?

The seven most common OTUs from the foliar fungal isolates collected from JRBP in 2015–2017 (i.e., the focal OTUs) comprised 66% and 77% of the isolates from native perennial and non-native annual grasses, respectively, across the density gradients (Table 2). None of the focal OTUs significantly increased in relative abundance with native perennial or non-native annual grass density (Tables S4–S5). The unidentified *Pyrenophora* species, however, tended to increase with higher grass density of either host group in the manipulated experiment (Fig. S5, Table S5).

**Table 2.**
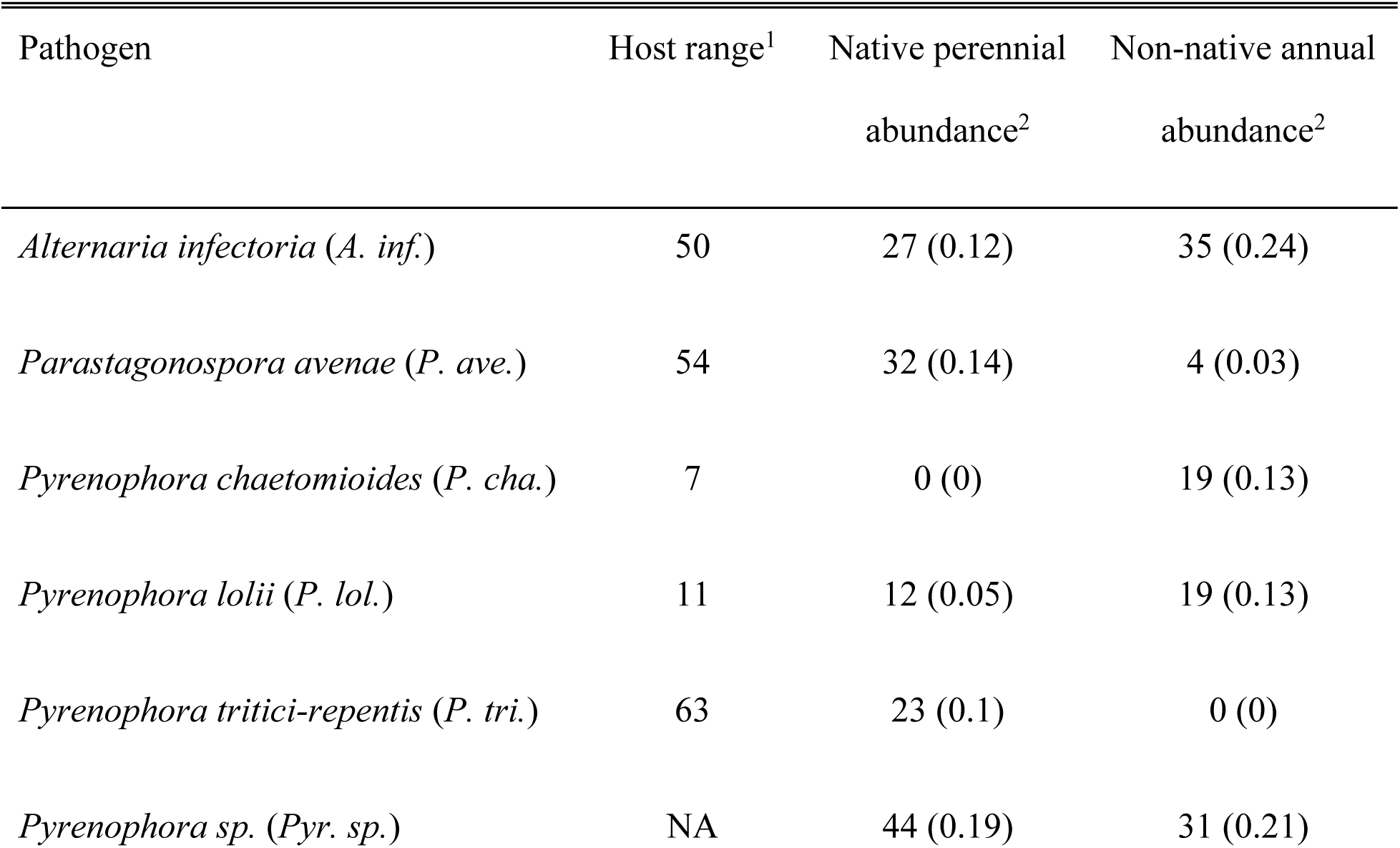
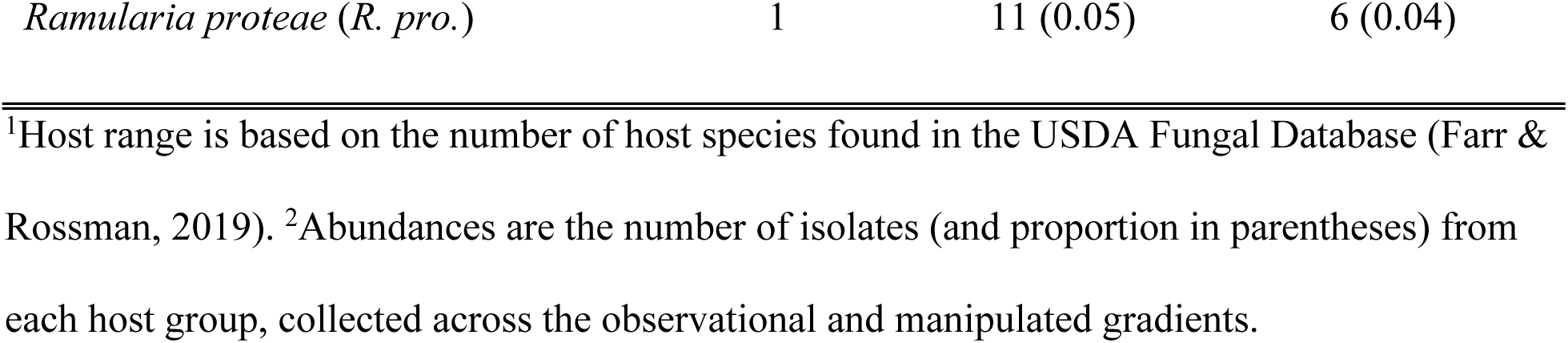
Pathogen species assigned to the focal OTUs.

We calculated the predicted change in relative abundance of each pathogen on each host group with the addition of 50 native perennial grasses m^-2^ (Fig. 3A–B) or 5000 non-native annual grasses m^-2^ (Fig. 3C–D) to bare plots. Such increases in density exceed those recorded in the observational study (Fig. S2A), but they still had small predicted impacts on the relative abundances of most pathogens (Fig. 3). Although not statistically significant, predicted *P. lolii* relative abundance decreased with 50 additional native perennial grasses (Fig. 3A–B), predicted relative abundance of the unidentified *Pyrenophora* species increased with 5000 additional non-native annual grasses (Fig. 3C–D), and predicted *P. chaetomioides* relative abundance on non-native annuals decreased with 5000 additional non-native annual grasses in the manipulated experiment (Fig. 3D).

**Figure 3.**
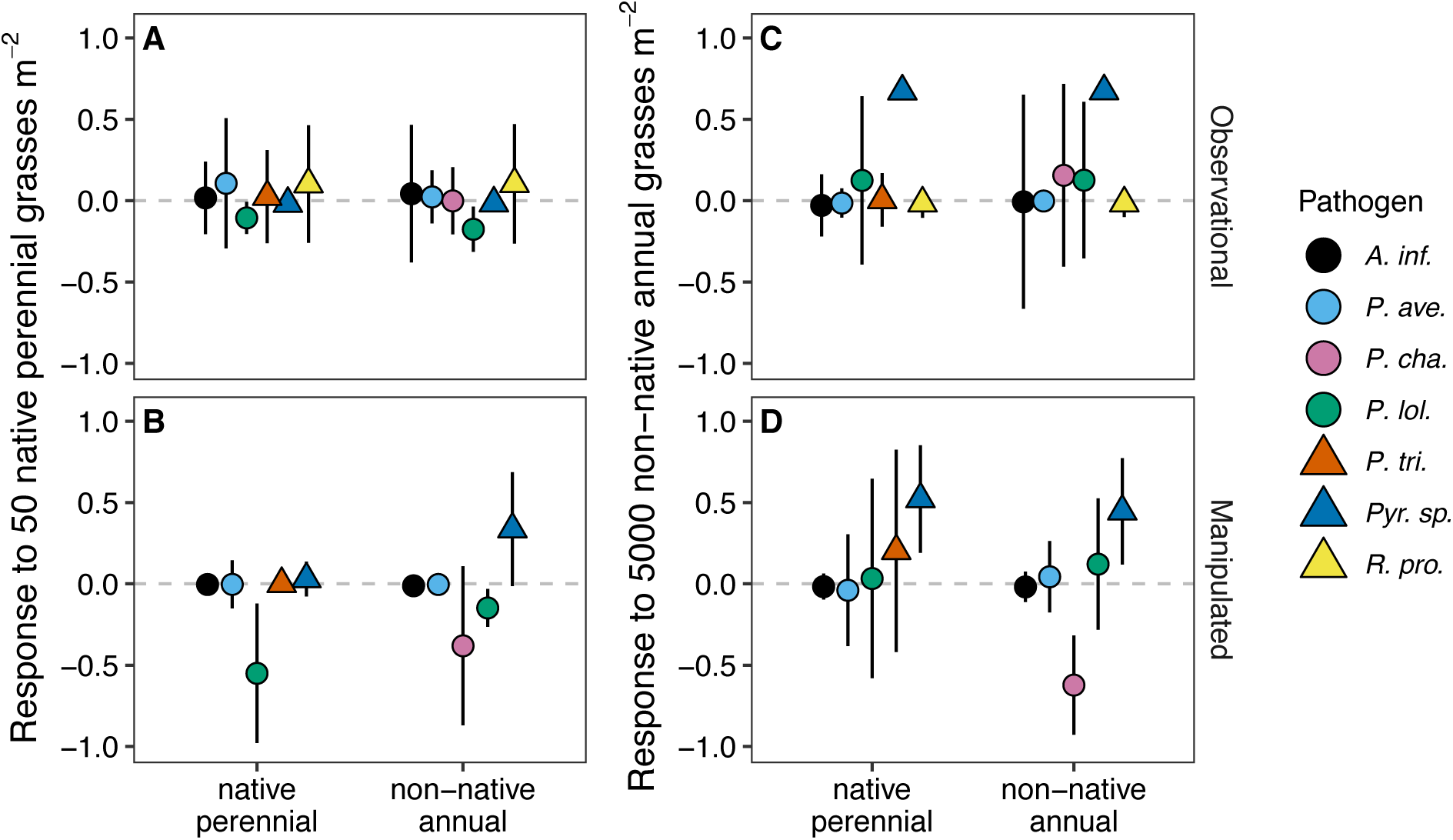
The average predicted effect (± 1SE) of adding (A–B) 50 native perennial grass individuals m^-2^ or (C–D) 5000 non-native annual grass individuals m^-2^ to a bare plot on the relative abundance of each of the seven focal OTUs on each host group (x-axes) based on regressions fit to the (A and C) observational and (B and D) manipulated studies (Tables S4–S5). Pathogen abbreviations are in Table 2.

### How do native perennial and non-native annual grass densities affect disease severity?

Disease severity, evaluated as the proportion of leaf surface area with lesions, was generally low across both host groups and grass density ranges (Fig. 4) and did not significantly change with grass density (Tables S6 and S7). The proportion of leaves with lesions also did not significantly change with native perennial or non-native annual grass density (Table S6 and S7), but tended to increase on native perennial hosts with higher densities of either host group in the observational study (Fig. 4A, C). However, higher disease severity at lower grass densities later in the season dampened these positive trends (Fig. S6). Mirroring results from the pooled dataset (Fig. 2), the proportion of leaves with lesions was 70% higher on native perennial grasses than non-native annual grasses in the observational study (*P* < 0.001, Table S6) and 46% higher in the manipulated experiment (*P* < 0.001, Table S7).

**Figure 4.**
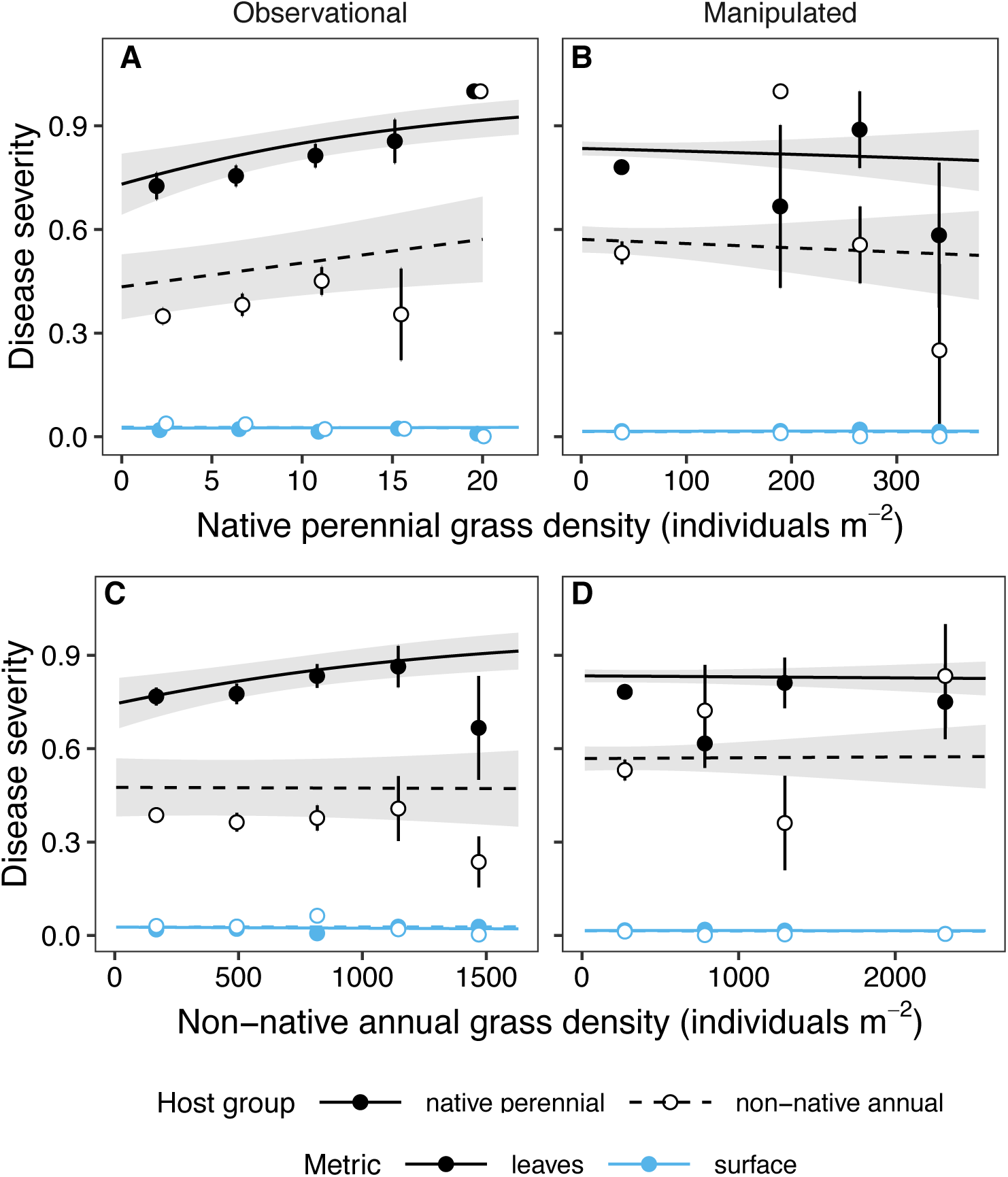
The effect of (A–B) native perennial and (C–D) non-native annual grass density on disease severity of native perennial and non-native annual hosts in the (A and C) observational study and (B and D) manipulated experiment. Disease severity was quantified as the proportion of leaves with lesions (“leaves”) or the average proportion of leaf surface area with lesions (“surface”) per plant. Density ranges were divided into five evenly spaced intervals and points representing the average disease severity within that interval (mean ± 1SE) are plotted at the midpoint. Points and error bars were nudged horizontally to reduce overlap. Lines and shaded regions represent linear regression fits (mean ± 1SE, Tables S6–S7). Shaded regions for surface are too small to visualize.

## Discussion

To evaluate the effects of host competence and density on pathogen community composition and disease severity, we isolated foliar fungi and quantified lesions on native perennial and non-native annual grasses in a California grassland across observational and manipulated densities of each group. We hypothesized that native perennials would host more specialized pathogens because of longer evolutionary histories with local pathogens, that non-native annuals would experience higher disease severity because of lower allocation to defenses, and that the two groups would host overlapping pathogen communities because of the long residence time of non-native annuals. We found the opposite for the first two hypotheses: non-native annuals hosted more specialized pathogens and native perennials experienced higher disease severity. The two groups hosted statistically distinct pathogen communities, but shared generalist pathogens. We also hypothesized that increases in the density of either group would induce shifts in pathogen community composition and increase disease severity, but the densities of both groups had no significant effects. Our findings suggest that even with variation in competence between two host groups, their densities do not necessarily influence components of disease that can affect risk, at least at the 1-m^2^ plot scale and over the three-year time span of our study.

### How does pathogen community composition differ between native perennial and non-native annual grass hosts?

Plant species vary in susceptibility to different pathogens (Barrett et al., 2009; Gilbert & Webb, 2007), in part due to life history (Cronin et al., 2010) and evolutionary history (I. M. Parker & Gilbert, 2004). Accordingly, native perennial and non-native annual grasses had distinct foliar fungal pathogen communities. These differences were driven by 59 pathogens that were only isolated from one host group and variation in isolation frequency of 24 pathogens shared between the groups. Hosts are frequently infected by multiple pathogens and, as demonstrated by our results and others, many pathogens can circulate among different hosts within the same community (Dobson et al., 2008; Griffiths et al., 2011; Malpica et al., 2006). Our study is unique, however, in seeking to understand how hosts that differ in life and evolutionary histories shape aboveground pathogen community composition (but see Seabloom, Borer, Lacroix, Mitchell, & Power, 2013). A pathogen community perspective demonstrated that high relative susceptibility of one group to one pathogen (e.g., native perennial grasses to *P. tritici-repentis*) can be balanced by high relative susceptibility of another group to another pathogen (e.g., non-native annual grasses to *P. chaetomioides*). This insight is likely to be general given that variation in evolutionary history also shapes soil microbial community composition (Kourtev, Ehrenfeld, & Häggblom, 2002; Lau & Suwa, 2016; Wolfe, Rodgers, Stinson, & Pringle, 2008) and cautions against conclusions about disease risk that focus on a single pathogen (Lloyd-Smith, 2013).

The enemy release hypothesis posits that native plants will experience greater disease pressure than non-native plants because non-native species will lose specialist enemies as they are transported to a new region and resident specialist enemies will be slow to attack non-native plants (Keane & Crawley, 2002). We isolated more unique OTUs from native perennials than non-native annuals, which supports this hypothesis, but we also found that the average host range of pathogens associated with non-native annuals was more specialized than that of pathogens associated with native perennials. The latter result should be interpreted with caution, however, because many of the pathogens did not have host range information available in the fungus database and some estimates of host range were inconsistent with our findings at JRBP (e.g., *R. proteae*). Nonetheless, the pathogens associated with non-native annual grasses may have a more specific host range for a few reasons. At least some of the pathogens that infect these grasses are globally distributed (Aboukhaddour, Cloutier, Lamari, & Strelkov, 2011; Stukenbrock, Banke, & McDonald, 2006), suggesting that specialists associated with non-native annual grasses in their native range may be present at JRBP. In addition, the non-native annual grass species were observed in JRBP or nearby counties as early as 1893 (JRBP, 2020), allowing time for pathogen adaptation to novel hosts (Hawkes, 2007; I. M. Parker & Gilbert, 2004).

### How does disease severity differ between native perennial and non-native annual grass hosts?

A higher proportion of native perennial leaves had lesions than non-native annual leaves. While this finding contradicts our expectation that non-native annuals would experience higher disease severity because of life history, it is consistent with multiple studies demonstrating higher disease severity on native than non-native plants (Chun, van Kleunen, & Dawson, 2010; Han, Dendy, Garrett, Fang, & Smith, 2008; Hawkes, Douglas, & Fitter, 2010; but see I. M. Parker & Gilbert, 2007). Native perennials may be more exposed to transmission and/or more susceptible to infection than non-native annuals, but, because they had equivalent and low proportions of leaf surface area with lesions, may possess equally effective resistance mechanisms. Differences in exposure may be partially explained by the long-lived life-history of native perennials and their role as long-term pathogen reservoirs (Thrall et al., 1993). Indeed, the difference in disease severity between non-native perennials and native annuals was greater in the observational study than the manipulated experiment, where plant communities had been recently assembled. In addition, non-native annual grasses may shed leaves with foliar fungal lesions (Vloutoglou & Kalogerakis, 2000) more frequently than native perennials, creating the appearance of lower disease severity.

### How do native perennial and non-native annual grass densities affect pathogen community composition?

Changes in density of native perennial and non-native annual grasses had limited effects on the relative abundance of foliar fungal pathogens. When we extrapolated our findings to a high density of grasses (50 and 5000 m^-2^ for native perennial and non-native annual, respectively), the predicted response of *P. chaetomioides* was inconsistent with the pathogen communities characterized with the larger dataset. *Pyrenophora chaetomioides* was predicted to decrease in relative abundance on non-native annuals with increasing density of non-native annuals, even though it was isolated frequently from two of the species in this group. It is possible that changes in plant community composition over the density gradient, such as an increase in *Bromus* spp. and a decrease in *Avena* spp., drove this pattern, suggesting that the densities of host genera or species may have contrasting effects on pathogen relative abundance.

Our results indicate that shifts in the densities of hosts that have similar life history strategies and local residence times do not necessarily shape the assembly of pathogen communities. While interpretations of biodiversity-disease risk relationships often invoke a strong role for host density (e.g., Young et al., 2017), pathogen communities may be more influenced by other factors, such as interspecific microbial interactions. For example, priority effects can influence the assembly of yeast communities in flower nectar (Peay, Belisle, & Fukami, 2012) and foliar fungal communities on grasses (Halliday, Umbanhowar, & Mitchell, 2017). One limitation to evaluating disease risk by particular pathogens in our study is that we lack data on the absence of infection. In addition, transmission events may occur at a scale greater than study plots, causing the plot-level density of grasses to be an inaccurate estimate of transmission pressure.

### How do native perennial and non-native annual grass densities affect disease severity?

While increases in grass density had no significant effect on disease severity, the proportion of native perennial leaves with lesions tended to increase over the observed density gradients. Despite a larger proportion of leaves being infected, the average proportion of leaf area with lesions was consistently low across grass densities, suggesting that grass densities may influence the number of times plants are exposed to pathogens, but not the potential for pathogens to grow on each plant. Deviations of our results from strong positive relationships between host density and disease severity (Mitchell et al., 2002; I. M. Parker et al., 2015) may be explained by plant community differences. High host diversity at JRBP may hinder foliar fungal pathogen adaptation to specific host defenses (Mundt, 2002), making pathogens less capable of exploiting locally-abundant hosts, and in turn less sensitive to changes in the density of any particular host group. Indeed, host percent cover in diverse old fields did not affect disease severity caused by aboveground pathogens (Schmidt et al., 2020). In addition, density effects may be transient, as exemplified by dampened density-disease severity relationships later in the growing season.

## Conclusions

This study of foliar fungal pathogen communities and resulting disease severity on native perennial and non-native annual grasses suggests that differences in life history or local residence time may contribute to disease risk through differences in competence, but not through changes in density. We could not parse out the independent effects of life history and local residence time, but previous studies of plant diseases suggest that both life history (Cronin et al., 2010) and evolutionary history (Gilbert & Webb, 2007; I. M. Parker et al., 2015) are strong drivers of competence. Our results are likely to be generally relevant, however, because non-native plants are more likely to be annuals and less likely to be perennials than native plants (Razanajatovo et al., 2016; Sutherland, 2004). These findings have implications for understanding the impacts of invasive species, which can cause species extinctions and shape ecological communities (Doherty, Glen, Nimmo, Ritchie, & Dickman, 2016; Pyšek et al., 2012). For example, when species initially invade a community, they can affect total host density, altering disease prevalence (Searle et al., 2016). The invasive species we evaluated, however, are well-established, suggesting that the expected impacts of invasive species on disease risk may be greater earlier in invasions. Our study demonstrates that host community composition can affect pathogen community composition and disease severity through variation in competence among hosts.

## Acknowledgements

The project was supported by the Jasper Ridge Kennedy endowment fund, the Stanford University Vice Provost for Undergraduate Education summer research fellowship for Biology undergraduates, the Bio-X summer research internship, and the Stanford University Raising Interest in Science and Engineering (RISE) summer internship program. We received research assistance and support from Nona Chiariello, Joe Wan, Phillippe Cohen, Teri Barry, Reuben Brandt, Joe Sertich, Johannah Farner, Steve Gomez, Bill Gomez, Stuart Koretz, Cary Tronson, Divya Ramani, Ryan Tabibi, Esther Liu, Sandya Kalavacherla, Vidya Raghvendra, Jason Zhou, Virginia Parra, Claudia Amadeo-Luyt, Guarika Duvar, and Elizabeth Wallace. AEK and SLF were supported by USDA award number 2017-67013-26870 as part of the joint USDA-NSF-NIH Ecology and Evolution of Infectious Diseases program. EAM was supported by an NSF Ecology and Evolution of Infectious Diseases grant (DEB-1518681), an NIH Maximizing Investigators’ Research Award (1R35GM133439-01), the Hellman Faculty Scholarship, and the Terman Award.

## Authors’ contributions

EAM and ERS designed the research, ERS and SCD conducted the field work and laboratory work, AEK conducted the analyses and wrote the first draft of the manuscript, all authors contributed to manuscript revisions and approved the final version.

## Data availability

The data and code will be available on the Github repository https://github.com/aekendig/invasion-pathogen-communities and archived with Zenodo.

