## Supplementary material for "Native perennial and non-native annual grasses shape pathogen community composition and disease severity in a California grassland": N/A

**Table S1.** Total abundance of grass species in the observational and manipulated studies.

| Study | Year | Grass species | Abundance | Host group |
|---|---|---|---|---|
| observational | 2015 | *Avena barbata* | 2114 | non-native annual |
| observational | 2015 | *Avena fatua* | 1865 | non-native annual |
| observational | 2015 | *Brachypodium distachyon* | 2801 | non-native annual |
| observational | 2015 | *Bromus diandrus* | 2429 | non-native annual |
| observational | 2015 | *Bromus hordeaceus* | 9112 | non-native annual |
| observational | 2015 | *Bromus sterilis* | 178 | non-native annual |
| observational | 2015 | *Elymus glaucus* | 72 | native perennial |
| observational | 2015 | *Festuca myuros* | 116 | non-native annual |
| observational | 2015 | *Festuca perennis* | 652 | other grass (non-native perennial) |
| observational | 2015 | *Phalaris aquatica* | 88 | other grass (non-native perennial) |
| observational | 2015 | *Stipa pulchra* | 127 | native perennial |
| observational | 2016 | *Avena* sp.[1] | 1076 | non-native annual |
| observational | 2016 | *Brachypodium distachyon* | 1533 | non-native annual |
| observational | 2016 | *Bromus diandrus* | 983 | non-native annual |
| observational | 2016 | *Bromus hordeaceus* | 3235 | non-native annual |
| observational | 2016 | *Elymus glaucus* | 70 | native perennial |
| observational | 2016 | *Stipa pulchra* | 84 | native perennial |
| observational | 2016 | *unidentified grass* | 160 | other grass |
| manipulated | 2016 | *Avena sp.*[1] | 1559 | non-native annual |
| manipulated | 2016 | *Brachypodium distachyon* | 5941 | non-native annual |
| manipulated | 2016 | *Bromus diandrus* | 854 | non-native annual |
| manipulated | 2016 | *Bromus hordeaceus* | 15076 | non-native annual |
| manipulated | 2016 | *Elymus glaucus* | 1325 | native perennial |
| manipulated | 2016 | *Festuca myuros* | 199 | non-native annual |
| manipulated | 2016 | *Gastridium phleoides* | 15 | non-native annual |
| manipulated | 2016 | *Stipa pulchra* | 286 | native perennial |

[1]Identified *Avena sp.* at JRBP include *A. barbata* and *A. fatua* (JRBP 2020)

**Table S2.** Number of isolates obtained from foliar fungal lesions for each grass species.

| Year | Host group | Grass species | Isolates |
|---|---|---|---|
| 2015 | native perennial | *Elymus glaucus* | 33 |
| 2015 | native perennial | *Stipa pulchra* | 58 |
| 2015 | non-native annual | *Avena barbata* | 29 |
| 2015 | non-native annual | *Avena fatua* | 12 |
| 2015 | non-native annual | *Bromus diandrus* | 23 |
| 2015 | non-native annual | *Bromus hordeaceus* | 11 |
| 2016 | native perennial | *Elymus glaucus* | 133 |
| 2016 | native perennial | *Stipa pulchra* | 128 |
| 2016 | non-native annual | *Avena barbata* | 96 |
| 2016 | non-native annual | *Bromus diandrus* | 83 |
| 2016 | non-native annual | *Bromus hordeaceus* | 63 |
| 2017 | native perennial | *Elymus glaucus* | 76 |
| 2017 | native perennial | *Stipa pulchra* | 106 |
| 2017 | non-native annual | *Avena barbata* | 85 |
| 2017 | non-native annual | *Bromus diandrus* | 14 |
| 2017 | non-native annual | *Bromus hordeaceus* | 11 |

**Table S3.** Summaries of the fixed effects of generalized linear regressions of disease severity, measured as the proportion of surface area with lesions ("surface") and the proportion of leaves with lesions ("leaves"), across all samples and years.

| Severity metric | Variable | Estimate | Std. error | z value | *P* |
|---|---|---|---|---|---|
| surface | intercept | -3.817 | 0.041 | -93.460 | **<0.001** |
| surface | host group | -0.004 | 0.049 | -0.090 | 0.931 |
| leaves | intercept | 1.243 | 0.432 | 2.881 | **0.004** |
| leaves | host group | -1.468 | 0.110 | -13.409 | **<0.001** |

Notes: The distributions of the response variables were beta (for "surface") and binomial (for "leaves")—both logit-link. Estimate and Std. error are in units of "change in the log-odds of disease severity for one unit increase in the variable". The intercept represents a native perennial host. *P*-values indicating statistical significance ($P < 0.05$) are in bold.

**Table S4.** AICc-based model-averaged estimates of the fixed effects of generalized linear regressions of pathogen relative abundance in the observational study for *Alternaria infectoria* (*A. inf.*), *Parastagonospora avenae* (*P. ave.*), *Pyrenophora chaetomioides* (*P. cha.*), *Pyrenophora lolii* (*P. lol.*), *Pyrenophora tritici-repentis* (*P. tri.*), *Pyrenophora sp.* (*Pyr. sp.*), and *Ramularia proteae* (*R. pro.*).

| Pathogen | Variable | Estimate | Std. error | z value | *P* | Importance |
|---|---|---|---|---|---|---|
| *A. inf.* | intercept | -2.140 | 0.902 | 2.372 | **0.018** | NA |
| *A. inf.* | host group | 1.771 | 0.542 | 3.267 | **0.001** | 1.00 |
| *A. inf.* | native perennial density | 0.016 | 0.160 | 0.103 | 0.918 | 0.28 |
| *A. inf.* | non-native annual density | -0.098 | 0.403 | 0.243 | 0.808 | 0.32 |
| *A. inf.* | other grass density | 0.551 | 0.575 | 0.958 | 0.338 | 0.74 |
| *A. inf.* | group:nat. per. density | 0.003 | 0.097 | 0.026 | 0.979 | 0.05 |
| *A. inf.* | group:non. ann. density | 0.095 | 0.378 | 0.253 | 0.801 | 0.12 |
| *A. inf.* | group:other density | -0.643 | 0.706 | 0.911 | 0.362 | 0.62 |
| *P. ave.* | intercept | -3.651 | 3.420 | 1.068 | 0.286 | NA |
| *P. ave.* | host group | -2.329 | 1.085 | 2.146 | 0.032 | 1.00 |
| *P. ave.* | native perennial density | 0.094 | 0.267 | 0.350 | 0.726 | 0.31 |
| *P. ave.* | non-native annual density | -0.063 | 0.272 | 0.230 | 0.818 | 0.28 |
| *P. ave.* | other grass density | -7.064 | 17.383 | 0.406 | 0.685 | 0.46 |
| *P. cha.* | intercept | -2.276 | 1.521 | 1.496 | 0.135 | NA |
| *P. cha.* | native perennial density | -0.007 | 0.214 | 0.033 | 0.973 | 0.19 |
| *P. cha.* | non-native annual density | 0.055 | 0.199 | 0.277 | 0.781 | 0.23 |
| *P. cha.* | other grass density | -1.690 | 7.310 | 0.231 | 0.817 | 0.44 |
| *P. lol.* | intercept | -2.454 | 0.529 | 4.643 | **<0.001** | NA |
| *P. lol.* | host group | 0.905 | 0.625 | 1.448 | 0.148 | 0.84 |
| *P. lol.* | native perennial density | -0.410 | 0.456 | 0.899 | 0.368 | 0.65 |
| *P. lol.* | non-native annual density | 0.046 | 0.186 | 0.248 | 0.804 | 0.32 |
| *P. lol.* | other grass density | 0.238 | 0.244 | 0.974 | 0.330 | 0.66 |
| *P. lol.* | group:nat. per. density | 0.094 | 0.332 | 0.284 | 0.776 | 0.17 |
| *P. lol.* | group:non. ann. density | -0.007 | 0.112 | 0.059 | 0.953 | 0.05 |
| *P. tri.* | intercept | -3.781 | 3.023 | 1.251 | 0.211 | NA |
| *P. tri.* | native perennial density | 0.054 | 0.292 | 0.185 | 0.853 | 0.21 |
| *P. tri.* | non-native annual density | 0.009 | 0.188 | 0.050 | 0.960 | 0.20 |
| *P. tri.* | other grass density | -0.080 | 0.338 | 0.238 | 0.812 | 0.28 |
| *Pyr. sp.* | intercept | -4.840 | 2.232 | 2.168 | **0.030** | NA |
| *Pyr. sp.* | host group | -0.114 | 0.606 | 0.188 | 0.851 | 0.24 |
| *Pyr. sp.* | native perennial density | -0.631 | 1.035 | 0.610 | 0.542 | 0.43 |
| *Pyr. sp.* | non-native annual density | 0.617 | 0.616 | 1.002 | 0.316 | 0.68 |
| *Pyr. sp.* | other grass density | -0.378 | 3.745 | 0.101 | 0.920 | 0.25 |
| *R. pro.* | intercept | -2.397 | 0.424 | 5.648 | **<0.001** | NA |
| *R. pro.* | host group | -0.061 | 0.312 | 0.196 | 0.845 | 0.27 |
| *R. pro.* | native perennial density | 0.076 | 0.212 | 0.360 | 0.719 | 0.33 |
| *R. pro.* | non-native annual density | -0.025 | 0.178 | 0.140 | 0.889 | 0.25 |
| *R. pro.* | other grass density | -0.039 | 0.231 | 0.169 | 0.866 | 0.25 |
| *R. pro.* | group:nat. per. density | 0.003 | 0.075 | 0.035 | 0.972 | 0.02 |

Notes: Variables may be missing because the sample size was too small (see Methods) or because they were not included in the 95% confidence set of the best-ranked models. The distributions of the response variables were binomial (logit-link). Estimate and Std. error are in units of "change in the log-odds of pathogen relative abundance for one unit increase in the variable". The importance is the sum of Akaike weights over all models in which the variable is present. The intercept represents a native perennial host in a study plot with average native perennial and non-native annual grass density. Density values were centered and scaled. *P*-values indicating statistical significance ($P < 0.05$) are in bold.

**Table S5.** AICc-based model-averaged estimates of the fixed effects of generalized linear regressions of pathogen relative abundance in the manipulated experiment for *Alternaria infectoria* (*A. inf.*), *Parastagonospora avenae* (*P. ave.*), *Pyrenophora chaetomioides* (*P. cha.*), *Pyrenophora lolii* (*P. lol.*), *Pyrenophora tritici-repentis* (*P. tri.*), and *Pyrenophora sp.* (*Pyr. sp.*)

| Pathogen | Variable | Estimate | Std. error | z value | *P* | Importance |
|---|---|---|---|---|---|---|
| *A. inf.* | intercept | -1.823 | 0.221 | 8.237 | **<0.001** | NA |
| *A. inf.* | host group | -0.011 | 0.599 | 0.018 | 0.986 | 0.30 |
| *A. inf.* | native perennial density | -0.046 | 0.157 | 0.293 | 0.769 | 0.31 |
| *A. inf.* | non-native annual density | -0.022 | 0.119 | 0.183 | 0.855 | 0.25 |
| *A. inf.* | group:nat. per. density | -0.205 | 1.579 | 0.130 | 0.897 | 0.03 |
| *P. ave.* | intercept | -2.562 | 1.227 | 2.088 | 0.037 | NA |
| *P. ave.* | host group | -0.913 | 1.883 | 0.485 | 0.628 | 0.85 |
| *P. ave.* | native perennial density | -0.038 | 0.183 | 0.209 | 0.835 | 0.31 |
| *P. ave.* | non-native annual density | -1.517 | 3.159 | 0.480 | 0.631 | 0.45 |
| *P. ave.* | group:nat. per. density | -0.348 | 3.377 | 0.103 | 0.918 | 0.05 |
| *P. ave.* | group:non. ann. density | 1.597 | 3.241 | 0.493 | 0.622 | 0.26 |
| *P. cha.* | intercept | -6.515 | 3.315 | 1.965 | **0.049** | NA |
| *P. cha.* | native perennial density | -8.661 | 10.152 | 0.853 | 0.394 | 0.57 |
| *P. cha.* | non-native annual density | -5.997 | 5.366 | 1.118 | 0.264 | 1.00 |
| *P. lol.* | intercept | -8.333 | 3.566 | 2.337 | **0.019** | NA |
| *P. lol.* | host group | -1.884 | 1.916 | 0.983 | 0.326 | 0.66 |
| *P. lol.* | native perennial density | -19.274 | 13.427 | 1.435 | 0.151 | 0.95 |
| *P. lol.* | non-native annual density | 0.067 | 0.246 | 0.272 | 0.785 | 0.29 |
| *P. tri.* | intercept | -7.614 | 4.294 | 1.773 | **0.076** | NA |
| *P. tri.* | native perennial density | 0.072 | 0.467 | 0.155 | 0.877 | 0.22 |
| *P. tri.* | non-native annual density | 0.310 | 6.301 | 0.049 | 0.961 | 0.21 |
| *Pyr. sp.* | intercept | -0.551 | 0.766 | 0.719 | 0.472 | NA |
| *Pyr. sp.* | host group | 1.114 | 2.071 | 0.538 | 0.591 | 0.63 |
| *Pyr. sp.* | native perennial density | 0.250 | 0.216 | 1.155 | 0.248 | 0.78 |
| *Pyr. sp.* | non-native annual density | 1.176 | 2.073 | 0.568 | 0.570 | 0.74 |
| *Pyr. sp.* | group:nat. per. density | 3.588 | 5.611 | 0.639 | 0.523 | 0.37 |
| *Pyr. sp.* | group:non. ann. density | -0.918 | 2.038 | 0.451 | 0.652 | 0.25 |

Notes: Variables may be missing because the sample size was too small (see Methods) or because they were not included in the 95% confidence set of the best-ranked models. The distributions of the response variables were binomial (logit-link). Estimate and Std. error are in units of “change in the log-odds of pathogen relative abundance for one unit increase in the variable”. The importance is the sum of Akaike weights over all models in which the variable is present. The intercept represents a native perennial host in a study plot with average native perennial and non-native annual grass density. Density values were centered and scaled. *P*-values indicating statistical significance ($P < 0.05$) are in bold.

**Table S6.** AICc-based model-averaged estimates (for “leaves”) and summary (for “surface”) of the fixed effects of generalized linear regressions of disease severity, measured as the proportion of surface area with lesions (“surface”) and the proportion of leaves with lesions (“leaves”), in the observational study.

| Severity metric | Variable | Estimate | Std. error | z value | *P* | Importance |
|---|---|---|---|---|---|---|
| surface | intercept | -3.653 | 0.059 | -62.150 | **<0.001** | NA |
| surface | host group | 0.073 | 0.065 | 1.110 | 0.266 | NA |
| surface | native perennial density | 0.019 | 0.042 | 0.460 | 0.647 | NA |
| surface | non-native annual density | -0.048 | 0.047 | -1.020 | 0.307 | NA |
| surface | other grass density | -0.060 | 0.067 | -0.910 | 0.365 | NA |
| surface | group:nat. per. density | -0.038 | 0.067 | -0.580 | 0.565 | NA |
| surface | group:non. ann. density | 0.054 | 0.069 | 0.770 | 0.439 | NA |
| surface | group:other density | 0.035 | 0.078 | 0.440 | 0.658 | NA |
| leaves | intercept | 1.438 | 0.359 | 4.009 | **<0.001** | NA |
| leaves | host group | -1.540 | 0.139 | 11.117 | **<0.001** | 1.00 |
| leaves | native perennial density | 0.322 | 0.177 | 1.812 | 0.070 | 0.94 |
| leaves | non-native annual density | 0.278 | 0.174 | 1.598 | 0.110 | 0.86 |
| leaves | other grass density | 0.077 | 0.117 | 0.653 | 0.514 | 0.60 |
| leaves | group:nat. per. density | -0.198 | 0.194 | 1.022 | 0.307 | 0.65 |
| leaves | group:non. ann. density | -0.281 | 0.195 | 1.444 | 0.149 | 0.80 |
| leaves | group:other density | -0.006 | 0.085 | 0.074 | 0.941 | 0.14 |

Notes: Variables may be missing because they were not included in the 95% confidence set of the best-ranked models. The distributions of the response variables were beta (for “surface”) and binomial (for “leaves”)—both logit-link. Estimate and Std. error are in units of “change in the log-odds of disease severity for one unit increase in the variable”. The importance is the sum of Akaike weights over all models in which the variable is present. The intercept represents a native perennial host in a study plot with average native perennial and non-native annual grass density. Density values were centered and scaled. *P*-values indicating statistical significance ($P < 0.05$) are in bold.

**Table S7.** AICc-based model-averaged estimates of the fixed effects of generalized linear regressions of disease severity, measured as the proportion of surface area with lesions ("surface") and the proportion of leaves with lesions ("leaves"), in the manipulated experiment.

| Severity metric | Variable | Estimate | Std. error | z value | *P* | Importance |
|---|---|---|---|---|---|---|
| surface | intercept | -4.121 | 0.052 | 78.620 | **<0.001** | NA |
| surface | host group | -0.090 | 0.084 | 1.070 | 0.285 | 0.71 |
| surface | native perennial density | 0.004 | 0.021 | 0.178 | 0.859 | 0.31 |
| surface | non-native annual density | -0.009 | 0.022 | 0.407 | 0.684 | 0.36 |
| surface | group:nat. per. density | -0.010 | 0.039 | 0.253 | 0.800 | 0.11 |
| surface | group:non. ann. density | -0.002 | 0.020 | 0.100 | 0.920 | 0.06 |
| leaves | intercept | 1.606 | 0.143 | 11.255 | **<0.001** | NA |
| leaves | host group | -1.329 | 0.203 | 6.562 | **<0.001** | 1.00 |
| leaves | native perennial density | -0.031 | 0.072 | 0.426 | 0.670 | 0.37 |
| leaves | non-native annual density | -0.009 | 0.060 | 0.156 | 0.876 | 0.31 |
| leaves | group:nat. per. density | 0.004 | 0.054 | 0.068 | 0.946 | 0.08 |
| leaves | group:non. ann. density | 0.014 | 0.076 | 0.186 | 0.852 | 0.08 |

Notes: Variables may be missing because they were not included in the 95% confidence set of the best-ranked models. The distributions of the response variables were beta (for "surface") and binomial (for "leaves")—both logit-link. Estimate and Std. error are in units of "change in the log-odds of pathogen relative abundance for one unit increase in the variable". The importance is the sum of Akaike weights over all models in which the variable is present. The intercept represents a native perennial host in a study plot with average native perennial and non-native annual grass density. Density values were centered and scaled. *P*-values indicating statistical significance ($P < 0.05$) are in bold.

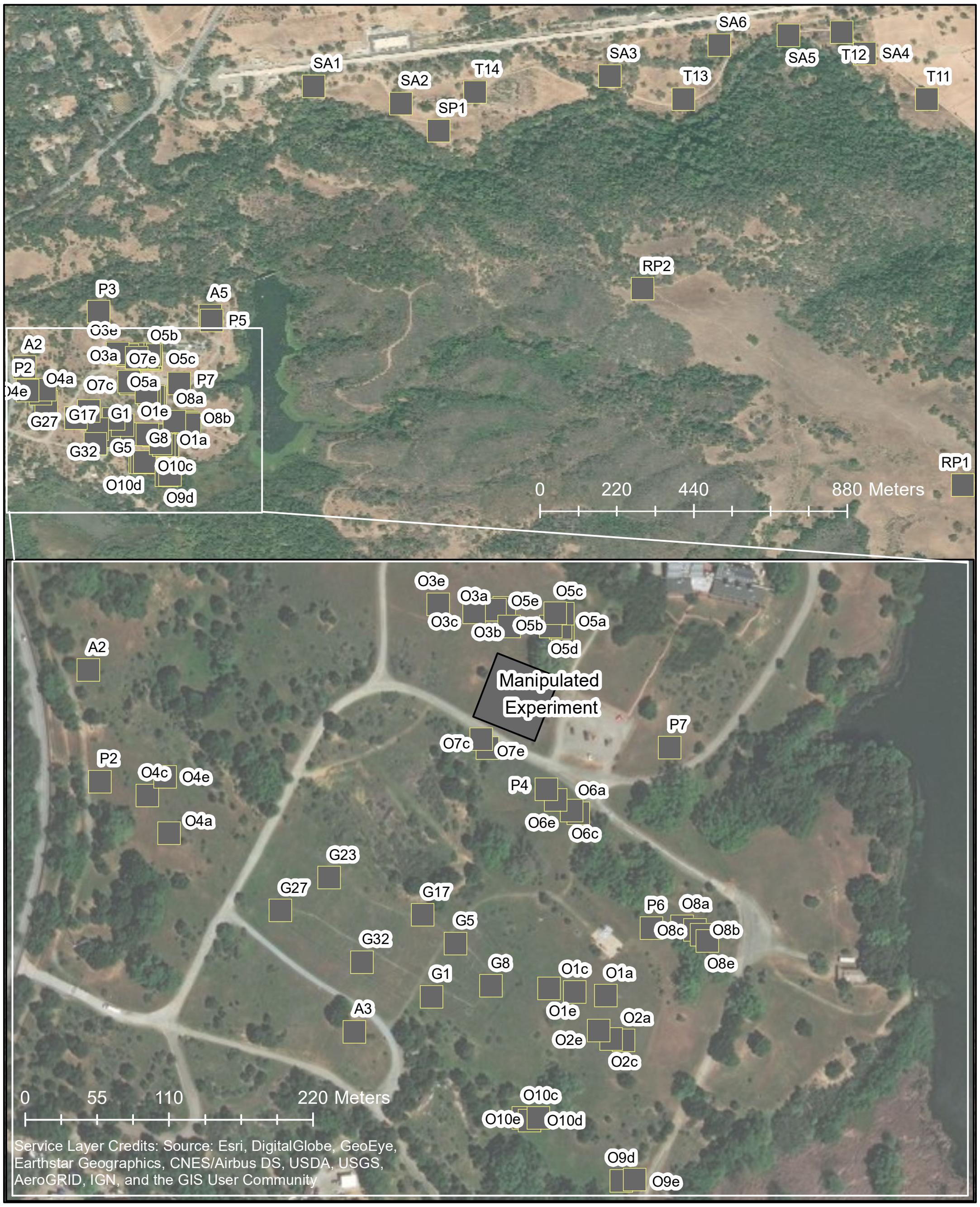


**Figure S1.** Map of sampling sites at Jasper Ridge Biological Preserve. The lower panel is a close-up view of a portion of the top panel. Letters indicate plot type: O: observational study, P: perennial dominated plots, A: annual dominated plots, G: control plots in the global change experiment, SA: sentinel plants in annual dominated plots, SP: sentinel plants in perennial dominated plots, T: transect of multiple plots, RP: ridge location of perennial dominated plots. The square labelled "Manipulated Experiment" contains all plots with manipulated density and additional ones used for a germinant study. For the observational study, the number indicates the transect and the lowercase letter indicates the plot within the transect.

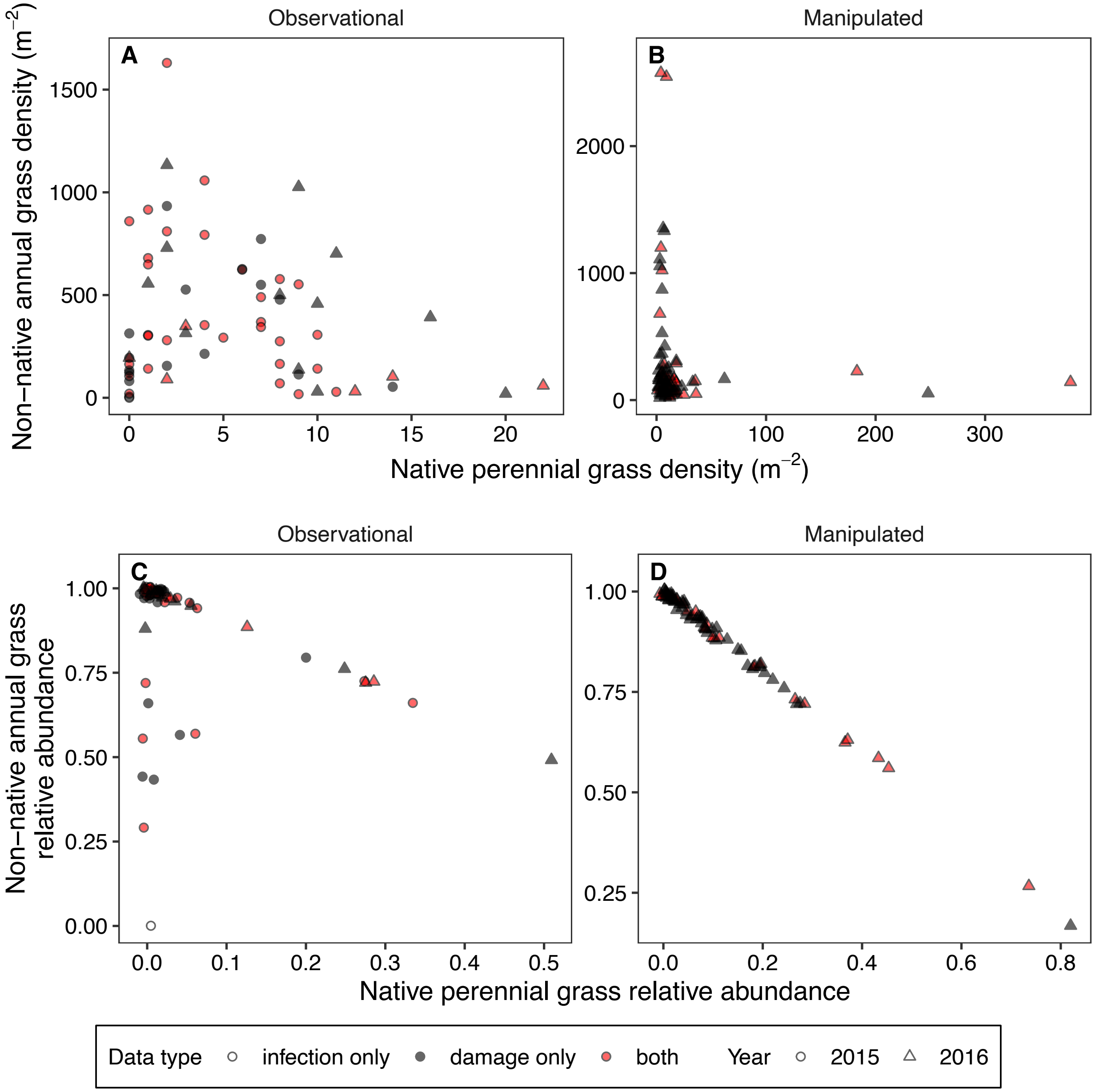


**Figure S2.** Native perennial and non-native annual grass density (A-B) and relative abundance (C-D) in the study plots. Each point represents a unique plot in a given experimental year (shape) and study (column). Data types collected from the plots include infection only (i.e., fungi isolated from lesions), damage only (i.e., the proportion of leaves and surface area with lesions), or both. Points in the relative abundance figures (C-D) are jittered away from center diagonal line for clarity.

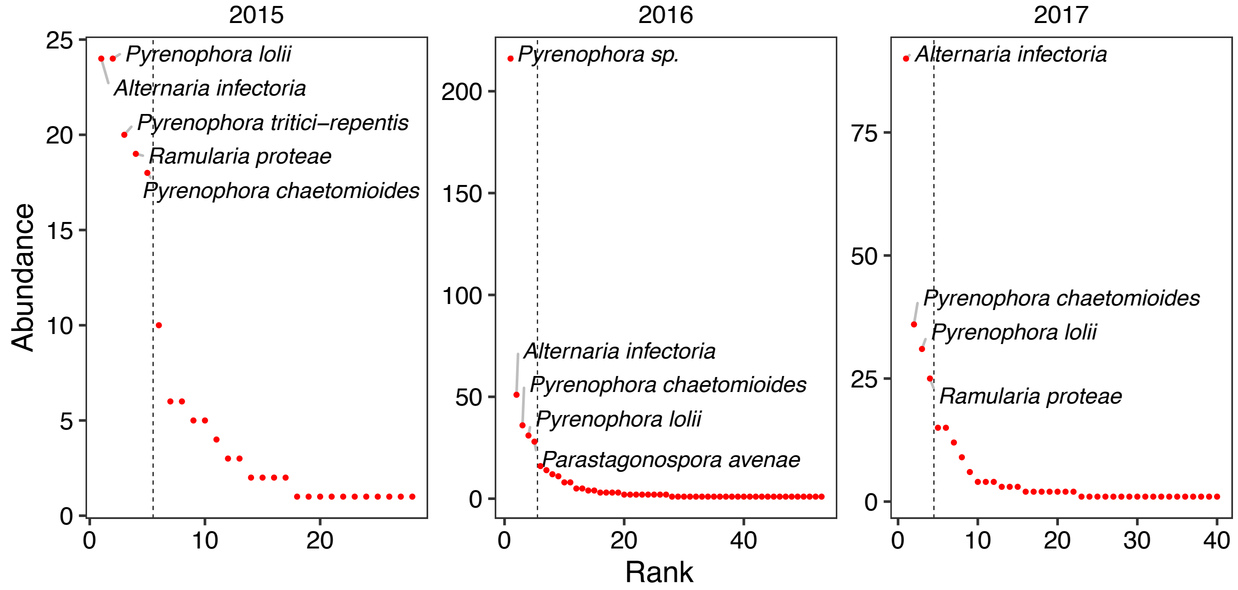


**Figure S3.** Relationship between abundance (number of isolates) and rank (determined by abundance) for fungal pathogen isolates from JRBP grasses in 2015, 2016, and 2017. Dashed vertical lines represent breaks used to determine the most common pathogens in each year (labelled points).

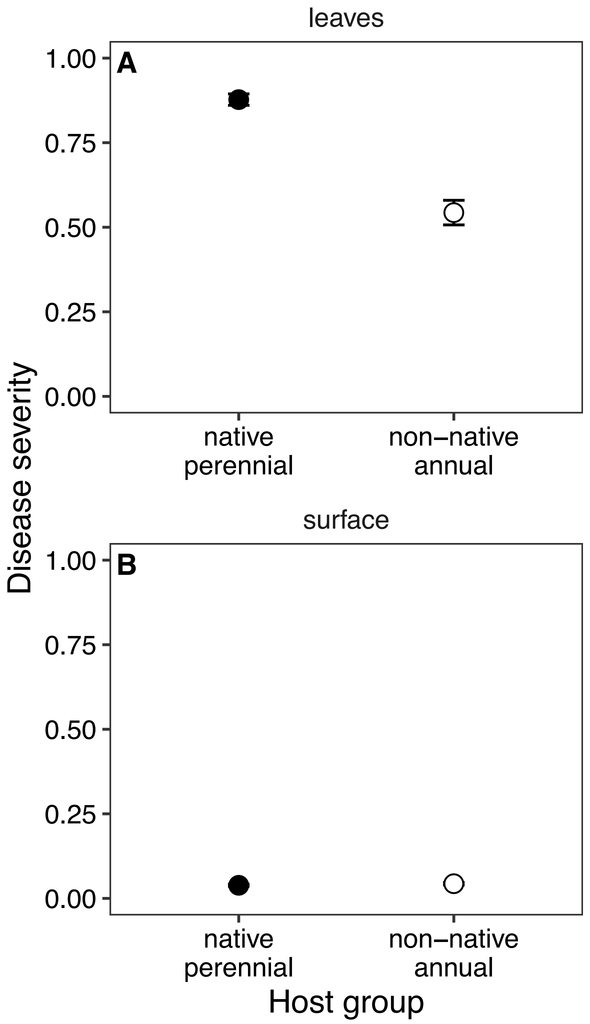


**Figure S4.** The average disease severity for each host group (model-estimated mean ± 1SE), quantified as (A) the proportion of leaves with lesions per plant and (B) the average proportion of infected leaf surface area with lesions per plant. This figure differs from Fig. 2 because the 2015 data from the observational study were collected in April 2015 rather than March 2015.

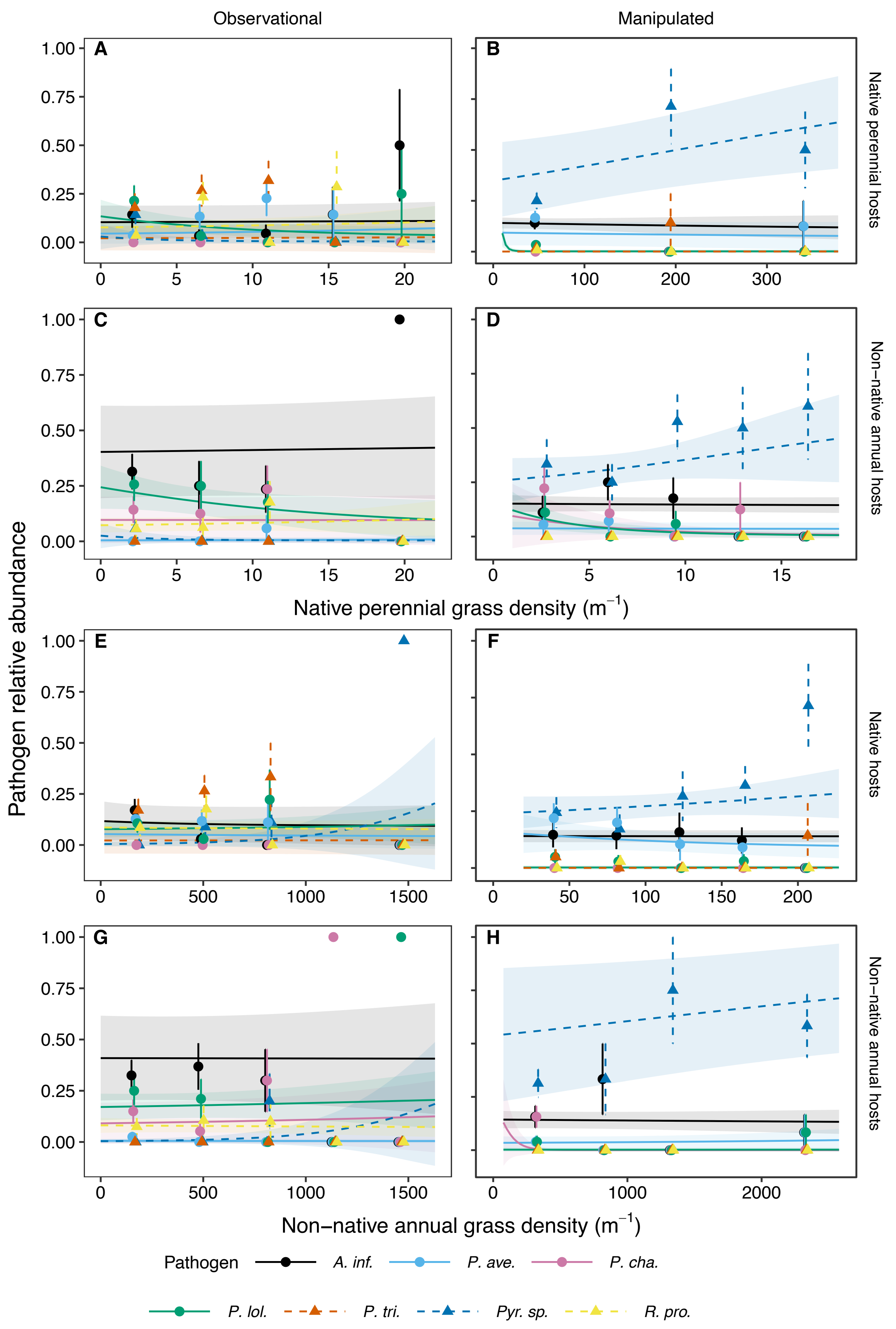

Observational
Manipulated
A
B
C
D
E
F
G
H
Native perennial hosts
Non−native annual hosts
Native hosts
Non−native annual hosts
Pathogen relative abundance
Native perennial grass density (m$^{-1}$)
Non−native annual grass density (m$^{-1}$)
Pathogen
A. inf.
P. ave.
P. cha.
P. lol.
P. tri.
Pyr. sp.
R. pro.

**Figure S5.** The effects of (A–D) native perennial and (E–H) non-native annual grass density on the relative abundances of the seven focal OTUs— *Alternaria infectoria* (*A. inf.*), *Parastagonospora avenae* (*P. ave.*), *Pyrenophora chaetomioides* (*P. cha.*), *Pyrenophora lolii* (*P. lol.*), *Pyrenophora tritici-repentis* (*P. tri.*), *Pyrenophora sp.* (*Pyr. sp.*), and *Ramularia proteae* (*R. pro.*). Columns represent the study type and rows represent the host group. Density ranges were divided into five evenly spaced intervals and points representing the average relative abundance of each pathogen within that interval (mean ± 1SE) are plotted at the midpoint. Points and error bars were nudged horizontally to reduce overlap. Lines and shaded regions represent linear regression fits (mean ± 1SE, Tables S4–S5).

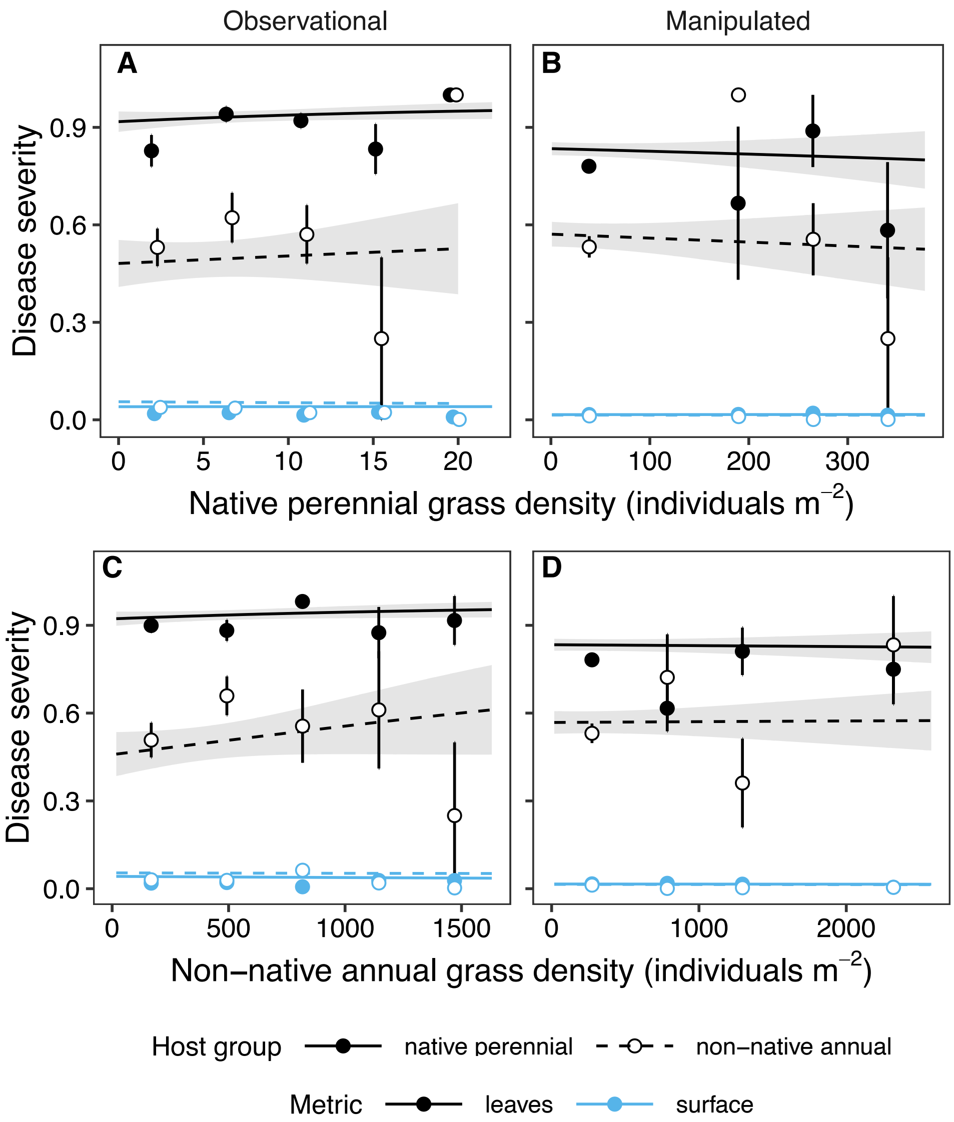


**Figure S6.** The effect of (A–B) native perennial and (C–D) non-native annual grass density on disease severity of native perennial and non-native annual hosts in the (A and C) observational study and (B and D) manipulated experiment. This figure differs from Fig. 4 because the 2015 data from the observational study were collected in April rather than March. Disease severity was quantified as the proportion of leaves with lesions ("leaves") or the average proportion of leaf surface area with lesions ("surface") per plant. Density ranges were divided into five evenly spaced intervals and points representing the average disease severity within that interval (mean ± 1SE) are plotted at the midpoint. Points and error bars were nudged horizontally to reduce overlap. Lines and shaded regions represent linear regression fits (mean ± 1SE, Tables S6–S7). Shaded regions for surface are too small to visualize.
